# Substrate-assisted Enzymatic Formation of Lysinoalanine in Duramycin

**DOI:** 10.1101/358382

**Authors:** Linna An, Dillon P. Cogan, Claudio D. Navo, Gonzalo Jiménez-Osés, Satish K. Nair, Wilfred A. van der Donk

**Author notes:** These authors contributed equally to this work (Linna An, Dillon P. Cogan).

## Abstract

Duramycin is a heavily post-translationally modified peptide that binds phosphatidylethanolamine. It has been investigated as an antibiotic, inhibitor of viral entry, therapeutic for cystic fibrosis, and tumor and vasculature imaging agent. Duramycin contains a β-hydroxylated Asp (Hya) and four macrocycles, including an essential lysinoalanine (Lal) crosslink. The mechanism of Lal formation is not known. We here show that Lal is installed stereospecifically by DurN via addition of Lys19 to a dehydroalanine. The structure of DurN reveals an unusual dimer with a new fold. Surprisingly, in the structure of duramycin bound to DurN, no residues of the enzyme are near the Lal. Instead, Hya15 of the substrate makes interactions with Lal suggesting it acts as a base to deprotonate Lys19 during catalysis. Biochemical data suggest that DurN preorganizes the reactive conformation of the substrate, such that the Hya15 of the substrate can serve as the catalytic base for Lal formation.

---

Duramycin and closely related compounds are ribosomally synthesized and post-translationally modified peptides (RiPPs) produced by various actinomycetes^1-5^ and marine symbionts^6^. Duramycin binds phosphatidylethanolamine (PE), a major structural phospholipid in mammalian and microbial cell membranes^7, 8^. Duramycin contains 19 amino acids, and five final posttranslational modifications (PTMs), including one lanthionine (Lan), two methyllanthionines (MeLan), an *erythro*-3-hydroxy-L-aspartic acid (Hya) resulting from the hydroxylation of Asp15, and one (2*S*,9*S*)-Lal (Figure 1). The carboxylate and hydroxyl groups of Hya15 make key interactions with the ethanolamine head group of PE, providing nanomolar affinity and high specificity^9^. As such, duramycin has been investigated as an antibiotic^1-3^, inhibitor of viral entry^10^, therapeutic for cystic fibrosis^11^, and tumor and vasculature imaging agent^12, 13^. Recently, divamide A, a close homolog of duramycin discovered from the symbiotic microbiome of small marine animals, was reported to display potent anti-HIV activity^6^.

**Figure 1:**
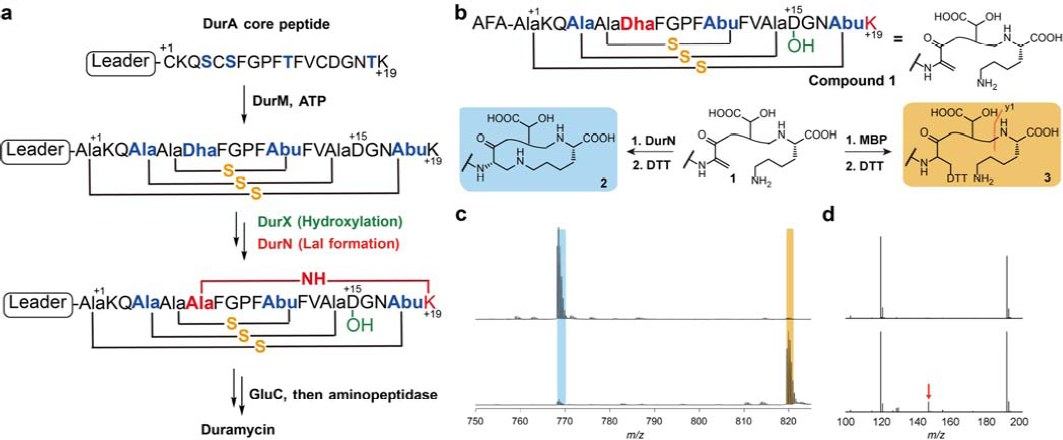
Duramycin biosynthesis. **a,** Co-expression of a minimal biosynthetic gene cassette and *in vitro* proteolytic removal of the LP produces duramycin. **b,** Peptide **1** formed by DurM and DurX is converted to **2** by urN *in vitro* (blue). Formation of the Lal is monitored by reaction with DTT, which provides an adduct (**3**) if Lal is not generated (yellow). The y1 fragment ion is marked with an orange line; **c,** Compound **1** was reacted with MBP-DurN (top) or MBP (bottom), treated with DTT, and analyzed by LC-MS (Supplementary Fig. 2). Expected and observed masses are listed in Supplementary Table 2. **d,** LC-MS/MS analysis of the products in panel c.

Previous studies have identified the biosynthetic genes (*dur*) that introduce the PTMs^14^. The precursor peptide DurA is synthesized with an N-terminal leader peptide (LP), which is removed after maturation of a C-terminal core peptide (CP) (Fig. 1a). Serine and threonine residues in the CP region of DurA are dehydrated by DurM to form dehydroalanine (Dha) and dehydrobutyrine (Dhb) residues. Subsequently, DurM catalyzes addition of cysteine thiols onto the β-carbon of Dha or Dhb to produce Lan or MeLan, respectively (Fig. 1a). The final two PTMs are hydroxylation of Asp15 by DurX to generate Hya15 and formation of (2*S*,9*S*)-Lal, putatively by DurN-catalyzed addition of Lys19 to Dha6 (Fig. 1a)^14^. A previous study demonstrated that duramycin lacking Lal does not have antimicrobial activity^14^, indicating that the Lal linkage is critical to set up the PE binding pocket. DurN and its orthologs have no sequence homology with any characterized proteins, its mechanism of Lal formation is not known, and previous attempts to reconstitute its activity *in vitro* were unsuccessful^15^.

In this study, we demonstrate DurN activity *in vitro* and present the co-crystal structures of the protein with a substrate analog and product. We compared Lal formation under enzymatic and nonenzymatic conditions, indicating that only DurN catalyzes stereospecific and diastereoselective Lal formation. Our data suggest that DurN promotes Lal formation via an unusual substrate-assisted mechanism.

## Results

### In vitro DurN activity

DurN was expressed in *Escherichia coli* as an N-terminal fusion with maltose binding protein (MBP-DurN; Supplementary Fig. 1). To prepare its putative substrate, the precursor peptide DurA was converted in *E. coli* to an intermediate containing Lan, MeLan, and Hya residues, but lacking Lal^14^. After purification, the majority of the leader peptide was removed proteolytically by endoproteinase Glu-C, yielding compound **1** with three N-terminal residues (AlaPheAla) originating from the leader peptide (Fig. 1a,b). Before testing the activity of DurN with this putative substrate peptide, we first generated the desired product by coexpression of DurA with DurM, DurX, and DurN in *E. coli*, followed by Glu-C treatment resulting in compound **2** containing all the PTMs found in duramycin (Fig. 1b)^14^. Because addition of Lys19 to Dha6 in peptide **1** to form the desired Lal does not lead to a change in mass nor a change in retention time, we adopted an indirect assay to follow Lal formation. Substrate **1** contains an electrophilic Dha, whereas product **2** does not. Hence, dithiothreitol (DTT) was used to monitor unreacted **1** by liquid chromatography-mass spectrometry (LC-MS) (Supplementary Fig. 2). After reacting **1** with MBP-DurN *in vitro* followed by DTT treatment, the DTT adduct **3** was not observed, whereas compound **3** was formed nearly quantitatively in control assays (Fig. 1c). The products were also analyzed by liquid chromatography-tandem mass spectrometry (LC-MS/MS), demonstrating that the y1 ion is observed for **1** and **3**, but not for the product of the MBP-DurN reaction because of the formation of the Lal ring (Fig. 1d; Supplementary Fig. 2b). Finally, as discussed in more detail in a later section, we hydrolyzed the product peptide of the MBP-DurN reaction and observed the presence of Lal by gas chromatography coupled to mass spectrometry (GC-MS). Thus, MBP-DurN catalyzes *in vitro* Lal formation in peptide **1** to form product **2**.

**Figure 2:**
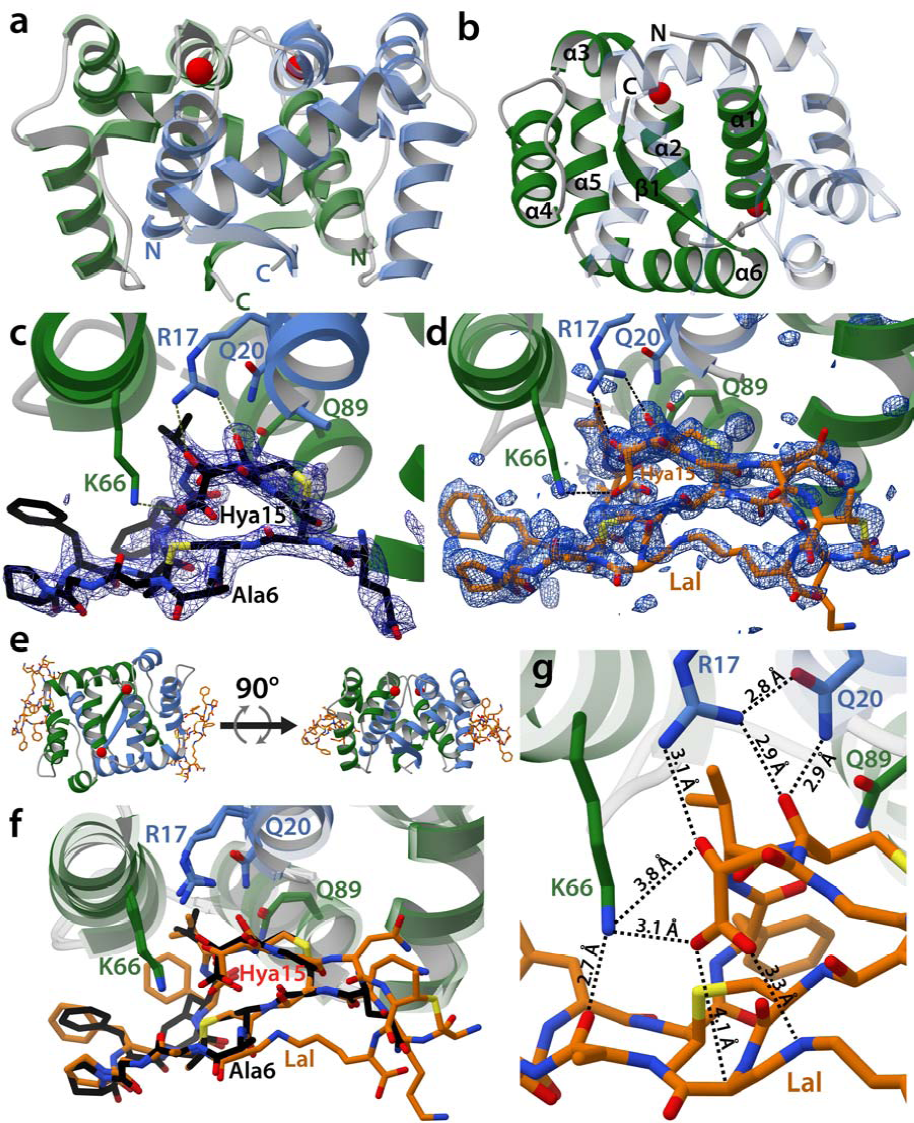
Structural analysis of DurN. **a,** Apo-DurN with the two monomers represented as green and blue ribbons. Two potassium ions are depicted as red spheres. **b,** Secondary structure mapped onto the DurN homodimer. For clarity, only the green monomer is labeled. **c,** DurN bound to **1-**Dha6Ala (black sticks). A difference Fourier map (F_o_ – Fc) contoured at 2.5σ is superimposed and shows electron density for portions of the ligand including one Lan, one MeLan, and Hya15. Critical catalytic DurN residues are represented as green and blue sticks. **d,** DurN bound to duramycin (orange sticks) with a difference Fourier map (F_o_ – F_c_) contoured at 2.5σ revealing occupancy for nearly the entire ligand. **e,** Two representations of the DurN homodimer featuring two molecules of duramycin related by a 90° rotational translation about the axis depicted by the black arrow. **f,** Superimposition of **1**-Dha6Ala and duramycin-bound structures of DurN. Phe7 in DurA is not modeled in the ligand structures (**c-d**, and **f**) because of insufficient electron density. **g,** Close-up view of the DurN-Hya15 interactions (interatomic distances are calculated averages between two ligand binding sites comprising chains C and G in PDB 6C0Y).

### The structure of DurN with a substrate analog

We removed the MBP tag from MBP-DurN (see Methods) and obtained crystal structures of DurN in complex with a substrate analog and in complex with duramycin. We generated an unreactive variant of **1** in which Dha6 (originating from Ser6 in DurA) was mutated to Ala (**1-**Dha6Ala, Supplementary Fig. 3a). The structure of DurN with **1-**Dha6Ala was solved to 1.90 Å resolution by single wavelength anomalous dispersion (SAD) methods using an iodide-soaked crystal. DurN also crystallized in the apo form and the structure was solved to 2.15 Å by molecular replacement using the SAD-derived coordinates (Fig. 2a). The overall structure of DurN consists of an interlaced homodimer where α-helix 1 of each monomer is engaged with residues from α-helices 2-6 in the adjacent monomer. The α-helices are followed by a single β-strand that forms an intermolecular, antiparallel interaction with its neighboring β-strand (Fig. 2a,b). A query against the PDB^16^ using the DALI server^17^ shows that DurN does not confidently resemble any known domain, indicating DurN contains a new protein fold and dimerization mode.

**Figure 3:**
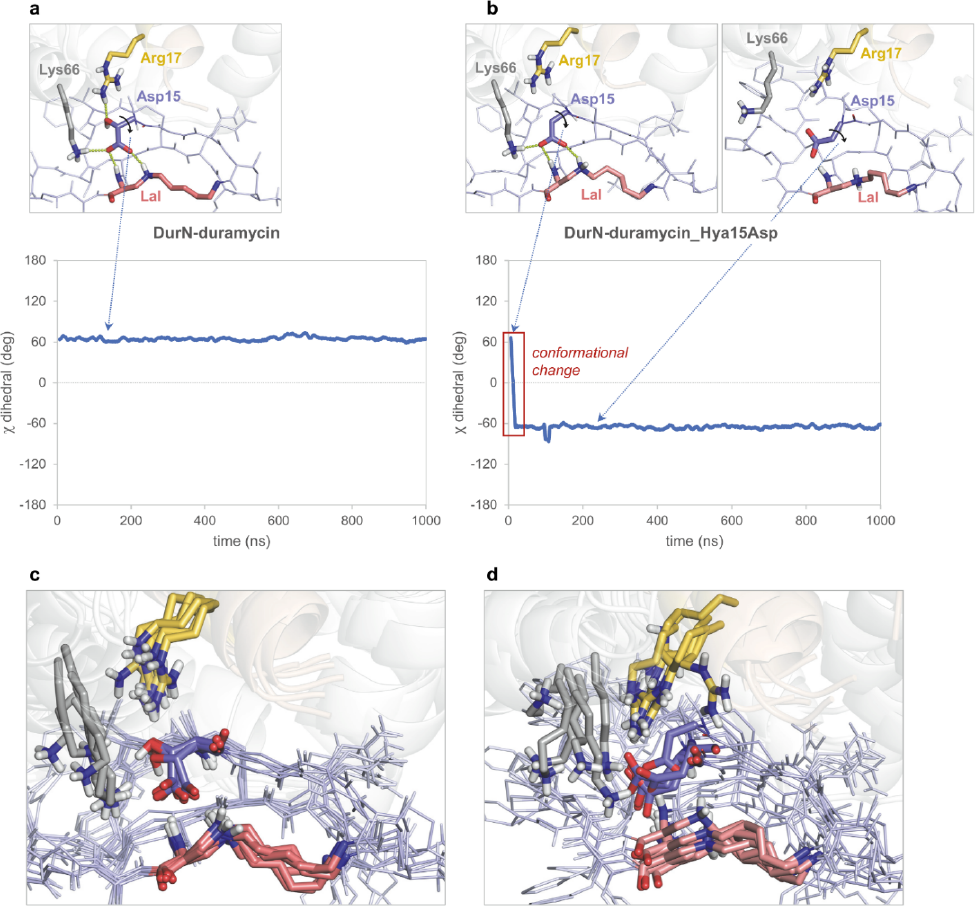
MD simulations indicating conformational changes of the Hya15Asp mutant compared to duramycin. The complex between DurN and duramycin was investigated through 1.0 μs MD simulations. Duramycin backbone and sidechains are shown as dark and light blue lines, respectively. Lysinoalanine (Lal), Hya/Asp15, Arg17 and Lys66 are shown as red, blue, yellow and grey sticks, respectively. DurN is shown as grey (chain A) and brown (chain B) transparent cartoons. Nonpolar hydrogens have been omitted for clarity. **a**, The hydrogen bond network between Arg17, Lys66 (both located in the enzyme DurN), Hya15 and Ala6 (both located in the substrate) observed in the crystallographic structures is maintained throughout the whole trajectory, as reflected by the conserved χ dihedral angle of Hya15 (carboxylate side chain) of *ca.* +60º. **b**, However, simply removing the β-hydroxy group in the Hya15Asp mutant disrupts this hydrogen bond network and compromises the integrity of the active site; the carboxylate side chain undergoes a 120º conformational twist and engages in electrostatic interactions with Arg17 and Lys66, trapping the catalytic base in a non-catalytic conformation as revealed by its χ dihedral angle. This computational observation rationalizes the lack of catalytic activity of DurN regarding Lal formation in the Hya15Asp mutant of **1**. Conformational ensembles for each complex show 5 superimposed representative snapshots derived from the trajectories and reveal a more rigid complex when native duramycin is bound to DurN (**c**), unlike the more flexible arrangement with the Hya15Asp mutant (**d**).

The homodimer contains two structural ions that are related by the pseudo two-fold symmetry axis of the DurN homodimer. Our data are consistent with potassium as the bound metal ions (Supplementary Figs. 4-5). DurN coordinates potassium at both sites via an octahedral array of oxygen ligands derived from amides in the backbone (Supplementary Fig. 4). In addition to potassium, we also considered the possibility of magnesium ions bound at the ion binding sites and carried out equivalent rounds of refinement in REFMAC5^18^ after modeling either potassium or magnesium ions independently. Whereas no positive signal was observed in the difference Fourier map (Fo – Fc) at 3σ for the corresponding potassium bound model, a positive difference signal was observed for the magnesium bound model (Supplementary Fig. 4c). Additional support for structural potassium ions was gathered by performing differential scanning fluorimetry (DSF), whereby DurN melting curves were obtained in the presence of various potassium or magnesium ion concentrations. While potassium ions conferred an increase in thermal stability of DurN as a function of ion concentration, magnesium ions had no effect on DurN thermal stability (Supplementary Fig. 4a,b). Molecular dynamics (MD) simulations also favor a structural potassium ion over magnesium (Supplementary Fig. 5).

**Figure 4:**
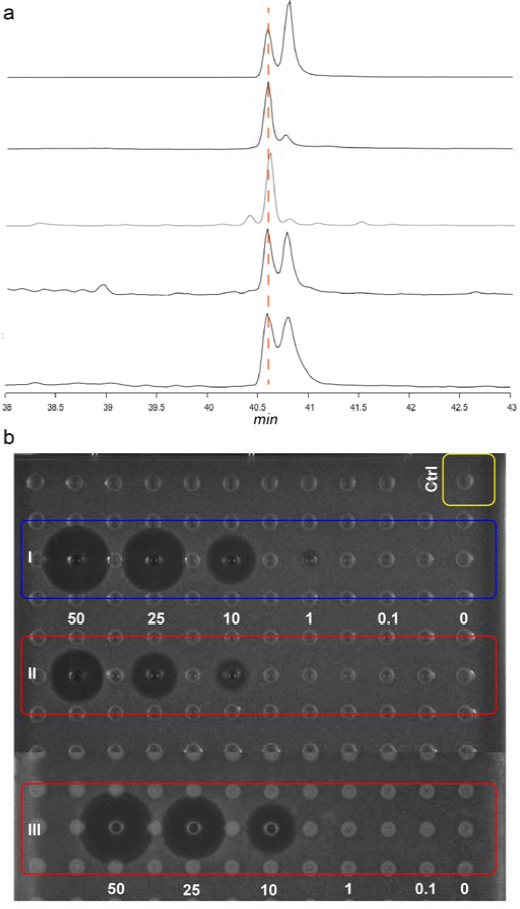
GC-MS analysis of derivatized lysinoalanine (Lal) and antimicrobial assay of duramycin and its analogs. **a,** commercial Lal standard that is a mixture of diastereomers (top trace), Lal obtained by hydrolysis of authentic duramycin (second trace), Lal from enzymatically formed **2** (third trace), Lal from non-enzymatically formed **2** (fourth trace), and Lal from hydrolyzed deoxyduramycin made in *E. coli* (bottom trace). **b,** Agar diffusion growth inhibition assay for duramycin generated by DurN *in vitro* (I), chemically generated **2** (II), and duramycin from the native producer (III). Concentrations are in μM, and the indicator strain is *Bacillus subtilis* ATCC 6633. Ctrl, negative control containing buffer and the proteases used to remove the leader peptide.

Analysis of simulated annealing difference Fourier maps (Fo – Fc) of the apo-structure and the structure with the bound substrate analog revealed electron density for **1-**Dha6Ala from Gln3 to Gly16 (Fig. 2c), with the reactive Lys19 invisible due to disorder. Only one of the two symmetric binding sites of the homodimer is occupied by ligand because of occlusion of the other site by residues from the N-terminus of a different protomer. Recognition of the ligand is mediated by residues from α-helix 1 of one polypeptide and α-helices 4-6 of the second monomer. Surprisingly, the ligand bound to the periphery of DurN and Ala6 (corresponding to the Dha6 involved in Lal formation) is far removed from any residues of the enzyme (Fig. 2c).

### The structure of DurN with duramycin

The observed binding pose of **1**-Dha6Ala does not provide an obvious mechanism of catalysis. We therefore determined the 1.66 Å resolution structure of DurN bound to duramycin. This structure revealed additional electron density in both binding sites of the DurN dimer, enabling modeling of the entire duramycin, including the Lal linkage (Fig. 2d,e). Globally, the two ligand-bound structures are highly similar with an RMSD of 0.45 Å over 1400 atoms (including ligands) (Fig. 2f). The duramycin-bound structure reaffirms that DurN does not provide any residues that could catalyze the Michael-type addition and suggests that DurN may mediate Lal formation by serving as a molecular scaffold to bring Dha6 and Lys19 in close proximity. Notably, a bidentate hydrogen-bond is observed between the guanidine of Arg17, the Hya15 hydroxyl group, and an amide backbone oxygen of Cys14 of duramycin (Fig. 2g). The Gln20 and Gln89 side chains of DurN further extend the hydrogen-bond network, and an electrostatic interaction is also observed between the carboxylate of Hya15 and Lys66 (Fig. 2g). These interactions position one of the carboxylate oxygens of Hya15 at 3.3 Å from the Lal secondary amine nitrogen. Thus, it appears that DurN holds the Hya15 carboxylate of the substrate in a conformation that is poised to activate Lys19 (Scheme 1). The other Hya15 carboxylate oxygen is 4.1 Å from the α-carbon of Ala6. This suggests that it would shield the *Si* face of the reactive Dha6, thus promoting protonation by solvent of the enolate formed upon the addition of Lys19 from the more accessible *Re* face, resulting in the correct (2*S*,9*S*)-Lal diastereomer (Scheme 1). The structures do not provide any evidence for activation of the carbonyl oxygen of Dha6. In fact, the Ψ angle in Ala6 in duramycin is close to zero, very likely due to electrostatic repulsion between the carboxylate of Hya15 and the amide carbonyl of Ala6, which is directed towards the solvent. This strongly suggests that the reacting Dha6 is fixed in an *s-cis* conformation prior to cyclization, which has been found to be the more reactive form in other Michael-type additions occurring in lanthipeptide biosynthesis^19^.

**Scheme 1:**
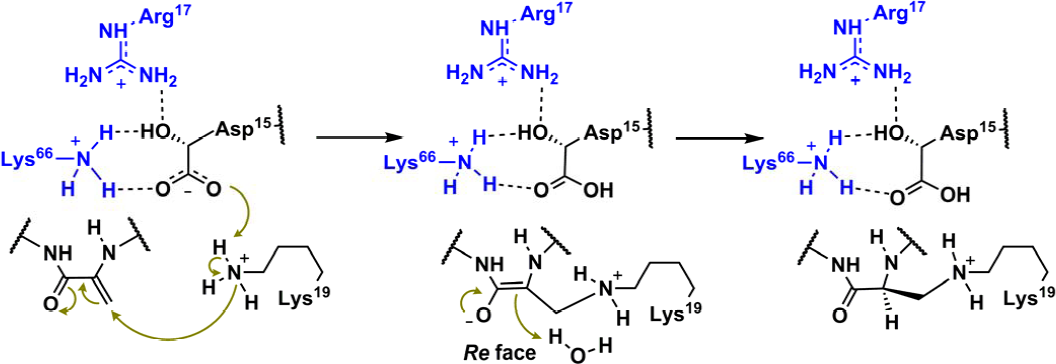
Proposed substrate assisted Lal formation catalyzed by DurN. Features in blue correspond to DurN and those in black correspond to its substrate/product.

### Substrate-assisted Lal formation

The conformation of duramycin in the co-crystal structure is more constrained than the solution structure of its close analog cinnamycin bound to lysophosphatidylethanolamine (PDB 2DDE). Indeed, molecular dynamics simulations demonstrate that the conformation of duramycin relaxes upon removal of DurN (Supplementary Fig. 6). Notably, in the absence of DurN, Hya15 moves away from the *Si* face of Dha6. Given the apparent importance of the hydroxy group of Hya15 for positioning its carboxylate, we also performed MD calculations on the complex in which this functionality was removed (i.e. Hya15 replaced by Asp). In the absence of the -hydroxy group, the carboxylate reorients to interact with Arg17 and Lys66, and thus the MD simulations predict it would no longer be available to act as a base (Fig. 3).

To probe their roles, MBP-DurN variants of conserved residues (Supplementary Fig. 7) were constructed and their activities determined with substrate **1**. Ala substitutions at positions involved in substrate binding (Arg17, Gln20, Gln89, and Lys66) resulted in inactive proteins, and the same outcome was observed for Arg18, Trp68, and Glu106 that are at the dimerization interface (Supplementary Fig. 8 & 9). To investigate the effect on affinity upon mutation of substrate binding residues, fluorescently labeled duramycin (**Dur-FL**; Supplementary Fig. 3) was used for fluorescence polarization (FP) studies. **Dur-FL** bound to MBP-DurN with a Kd of 417 ± 6 nM (Supplementary Fig. 10a). Unlabeled duramycin and substrate **1** were then used in competition FP experiments providing inhibition constants (Ki) of 2.72 ± 0.01 μM and 16.8 ± 0.5 μM, respectively (Supplementary Fig. 10b). MBP-tagged DurN variants R17A, Q20A, and Q89A did not bind Dur-FL, R17K bound very weakly, and K66A provided a Ki of 7.43 ± 0.08 μM (Supplementary Fig. 10b). Collectively, the mutational data suggest that the binding modes observed in the co-crystal structures are functionally important because variants of residues that engage the substrate in the crystal are compromised with regards to both catalysis and substrate binding.

We also tested variants of substrate **1**. To maintain the hydroxyl group of Hya but not its carboxylate, **1-**Hya15Ser (Supplementary Fig. 3c) was prepared in similar fashion as **1** except for using DurA-D15S as a precursor peptide. **1-**Hya15Ser was a poor substrate for MBP-DurN (Supplementary Fig. 11), even though competition FP experiments show that this variant peptide inhibits **Dur-FL** binding with a Ki of 10.2 ± 0.3 μM (Supplementary Fig. 10b). Another variant, **1-**Hya15Asp, was prepared by removing DurX from the *E. coli* co-expression system (Supplementary Fig. 3f). This variant was not processed by MBP-DurN *in vitro* (Supplementary Fig. 11), in agreement with the predictions of the MD simulations described above. These data suggest that both functional groups on the Hya15 side chain (i.e. β-hydroxyl and γ-carboxylate) must be present for substrate to be processed by DurN.

### Stereochemistry of enzymatic and non-enzymatic Lal formation

The observation that **1**-Hya15Asp was not a substrate for DurN *in vitro* was surprising because several studies have shown that deoxyduramycin/deoxycinnamycin can be obtained by co-expression of DurA/CinA with DurM/CinM and DurN/CinN^20, 21^. Thus, it appears that in bacteria, the absence of the hydroxy group on Asp15 is not preventing Lal formation. To further investigate this apparent contradiction, we obtained deoxyduramycin by co-expressing DurA with DurM and DurN in *E. coli* (Supplementary Fig. 3g). The product was treated with Glu-C to remove most of the leader peptide, and the three residues remaining from the leader peptide were removed using aminopeptidase. To probe whether duramycin and deoxyduramycin contained Lal, each peptide was hydrolyzed under acidic conditions to their constituent amino acids, which were derivatized as described previously (see Methods)^15^. GC-MS analysis indicated that authentic duramycin isolated from the producer organism contained predominantly the natural LL-Lal isomer (Fig. 4a), with a small amount of epimer that is likely formed during acid hydrolysis^6, 15^. However, deoxyduramycin formed in *E. coli* contained about equal amounts of two diastereomers. Peptide **1** was then exposed to alkaline conditions that were previously used to form the Lal non-enzymatically but for which the stereochemistry was not determined^15^. After removal of the leader peptide, hydrolysis and analysis by GC-MS, two diastereomers of Lal were detected in roughly equal amounts, which was also observed when using a commercial mixture of (2*R*,9*S*)-Lal and (2*S*,9*S*)-Lal standards (Fig. 4a). In contrast, product **2** obtained in vitro from DurN-catalyzed Lal formation again existed predominately of the (2*S*,9*S*)-Lal isomer. Collectively, these findings therefore suggest that Lal formation in *E. coli* when DurN is absent is non-stereoselective and may be nonenzymatic.

To explore the ramifications of the stereochemical information, we determined the antimicrobial activity of authentic duramycin, DurN-generated duramycin, and chemically generated duramycin by an agar diffusion assay against *Bacillus subtilis* ATCC 6633. These compounds would all contain the Hya15 and would only differ in the Lal. As expected based on the stereochemical analysis (Fig. 4a), the product of DurN catalysis showed a similar zone of growth inhibition as authentic duramycin, whereas nonenzymatically produced **2** showed a zone of inhibition roughly half the size (Fig. 4b).

### Conclusion

In this study, we provide the first *in vitro* demonstration that DurN catalyzes Lal formation, a step that had been recalcitrant in previous studies^6, 15^. By comparing enzymatic and non-enzymatic Lal generation, our data suggest that DurN greatly facilitates the reaction and controls the stereoselectivity. Furthermore, our data shows that the enzyme does not require the leader peptide for catalysis. DurN has a unique fold and dimeric architecture, and in the co-crystal structures, the Lal is entirely solvent exposed and the Hya15 of the substrate is positioned to potentially act as a base during catalysis to deprotonate Lys19 to initiate the Michael addition. Additionally, Hya15 appears to shield the *Si* face of the enolate intermediate, such that protonation will only occur from the *Re* face, resulting in the correct (2*S*,9*S*)-Lal product. Mutational studies show that substrates that lack either the hydroxyl or carboxylate groups of Hya15 are not processed by DurN, despite maintaining binding affinity, suggesting both of these groups on the substrate are indispensable for catalysis. Merging the biochemical findings with structural and computational insights points to a mechanism whereby the substrate employs its own Hya15 as a catalytic base to facilitate a Michael addition and stereospecific enolate protonation, resulting in Lal formation between Lys19 and Dha6.

## Acknowledgements

The authors thank Dr. L. Huo for providing plasmids, Dr. A. Vladimirovich Ulanov (UIUC Metabolomics Center) for technical assistance on GC-MS analysis, and K. Brister and the staff at LS-CAT at the Advanced Photon Source (Argonne National Laboratory) for their assistance with X-ray crystallography data acquisition. This work was supported by the National Institutes of Health (R37 GM 058822 to W.A.V.), and D.G.I. MINECO/FEDER (grants CTQ2015-70524-R and RYC-2013-14706 to G.J.O. and a predoctoral fellowship to C.D.N).

## Author Contributions

L.A. performed all biochemical assays, designed and analysed by L.A. and W.A.V., and D.P.C. and S.K.N. performed and interpreted all structural studies. C.D.N. and G.O. performed the computational analysis. L.A., D.P.C., S.K.N. and W.A.V. wrote the manuscript. L.A. and D.P.C. contributed equally to this study.

## Author Information

Atomic coordinates and structure factors for the reported crystal structure are deposited in the Protein Data Bank under accession codes 6C0G, 6C0H, and 6C0Y. Reprints and permissions information is available at www.nature.com/reprints. The authors declare no competing financial interests. Readers are welcome to comment on the online version of the paper. Correspondence and requests for materials should be addressed to S.K.N. or W.A.V.

## Methods

### General methods

Positive numbers are used for amino acids in the core peptide counting towards the C-terminus from the leader peptide cleavage site. Negative numbers are used counting backwards from the cleavage site. Polymerase chain reactions (PCRs) were carried out on an automated thermocycler (C1000^TM^, Bio-Rad). Gibson Assembly reaction solutions were made as reported^22^. DNA sequencing was performed by ACGT, Inc (Wheeling, IL). Matrix-assisted laser desorption/ionization time-of-flight mass spectrometry (MALDI-TOF MS) analyses were carried out on an UltrafleXtreme mass spectrometer (Bruker Daltonics). Samples were desalted using ZipTipC18 (Millipore) and spotted onto a MALDI target plate with a matrix solution containing 35 mg/mL 2,5-dihydroxybenzoic acid (DHB) in 3:2 MeCN/H2O with 0.1% trifluoroacetic acid (TFA). Peptides were desalted by C18 solid-phase extraction (SPE) and purified by reversed-phase high performance liquid chromatography (RP-HPLC) on a Shimadzu LC-20AP equipped with a Phenomenex Luna^^®^^ column (10 µm C18(2), 100 Å, 250 × 10 mm; part number: 00G-4253-N0) at a flow rate of 8 mL/min. An Agilent 1260 Infinity Quaternary LC System was employed for analytical HPLC with an Phenomenex Luna column (10 µm C18(2), 100 Å, 250 × 4.6 mm; part number: 00G-4253-E0) at a flow rate of 1 mL/min. Solvent A was 50 mM ammonium formate in H2O and solvent B was acetonitrile. A gradient of 5% solvent B to 60% solvent B over 30 min was used unless specified otherwise. Liquid chromatography electrospray ionization tandem mass spectrometry (LC/ESI-Q/TOF-MS/MS) was carried out using a Synapt ESI quadrupole TOF Mass Spectrometry System (Waters) equipped with an Acquity Ultra Performance Liquid Chromatography (UPLC) column (1.7 µm BEH C8, 130 Å, 1.0 × 100 mm, part number: 186002876). For all LC-MS or LC-MS/MS analysis, solvent A was water with 0.1% formic acid and solvent B was acetonitrile with 0.1% formic acid. An elution gradient of 15% solvent B to 60% solvent B over 20 min was used unless specified otherwise. For size-exclusion chromatography (SEC) studies, a HiLoad 16/60 column packed with Superdex 200 Prep Grade was used on an ÄKTA purifier eluting at 1.5 mL/min at 4 °C. GC-MS analysis was performed at the Roy J. Carver Metabolomics Center (UIUC) using an Agilent 7890 gas chromatograph with a ZB-1MS (25 m × 0.25 mm × 0.25 µm) column (Phenomenex).

### Materials

Oligonucleotides were obtained from Integrated DNA Technologies. Restriction endonucleases, DNA polymerases, and T4 DNA ligase were purchased from New England Biolabs. Media components were obtained from Difco Laboratories. Lysinoalanineꞏ2HCl was purchased from Bachem. Other chemicals were obtained from Sigma Aldrich or Fisher Scientific.

### Strains and plasmids

*E. coli* DH5α was used as host for cloning and plasmid propagation and *E. coli* BL21 (DE3) as host for expression of proteins/peptides. pRSFDuet-1, pACYCDuet-1, and pETDuet-1 were obtained from Novagen. The pET His6 small ubiquitin-like modifier (Sumo) tobacco etch virus (TEV) protease LIC cloning vector (2S-T) was a gift from Scott Gradia (Addgene plasmid #29711). Procedures for construction of all plasmids and preparation of all proteins can be found in Supplementary Information 1.

### Production and purification of 1, 2, 1-Dha6Ala, 1-Hya15Ser, 1-Hya15Asp, and deoxyduramycin

*E. coli* BL21(DE3) cells were transformed with pRSFDuet/His6-DurA/DurM and pACYCDuet/DurX to prepare compound **1** or with pRSFDuet/His6-DurA/DurM and pACYCDuet/DurN/DurX to prepare compound **2**. Cells were transformed with pRSFDuet/His6-DurA-Dha6Ala/DurM and pACYCDuet/DurX to prepare **1-**Dha6Ala, with pRSFDuet/His6-DurA-Asp15Ser/DurM and pACYCDuet/DurX to prepare **1-**Hya15Ser, or with pRSFDuet/His6-DurA/DurM to prepare **1-**Hya15Asp. Transformants were plated on a LB agar plate containing 50 mg/L kanamycin and 50 mg/L chloramphenicol. *E. coli* BL21 (DE3) cells were transformed with pRSFDuet/His6-DurA/DurM, pMAL-DurN to prepare deoxyduramycin and the transformants were plated on a LB agar plate containing 50 mg/L kanamycin and 50 mg/mL ampicillin. The following procedures are the same for all preparations. A single colony was picked and grown in 5 mL of LB at 37 °C for 12 h containing antibiotics as described above, and the resulting culture was used to inoculate 1 L of LB containing the same antibiotics. When the OD600 reached 0.75, the cultures were cooled on ice for 30 min, and then IPTG was added to 1.0 mM. The cells were cultured at 18 °C for another 18 h before harvesting by centrifugation at 8,000 × g for 10 min. Cell pellets were resuspended at room temperature in LanA buffer 1 (6 M guanidine hydrochloride, 500 mM NaCl, 0.5 mM imidazole, 20 mM NaH2PO4, pH 7.5 at 25 °C) followed by sonication. The lysates were centrifuged at 18,000 ×*g* for 30 min and supernatants were passed through 0.45-µm syringe filters. His-tagged modified peptides were purified by immobilized metal affinity chromatography (IMAC) loaded with nickel as previously described^13^. The fractions with the desired peptide were desalted by preparative RP-HPLC using an Agilent Bond Elut C18 column (3 mL). The desalted peptides were lyophilized, dissolved in 50 mM Tris pH 7.5 buffer, and digested by endoproteinase GluC at 20:1 (w/w). The digested reaction mixtures were purified by reversed-phase prep-HPLC (see General Methods).

To prepare deoxyduramycin, the peptide was digested with aminopeptidase (0.1 U per µg of peptide) at pH 8.5 at room temperature overnight and purified by analytical HPLC (General Methods).

### Dithiothreitol (DTT) assay to detect uncyclized Dha

DTT was dissolved in water at 1 M then aliquoted and stored at −20 °C for up to two weeks. DTT solutions were added to the peptide solutions to give a final concentration of 250 mM, and pH was adjusted (if necessary) to 7.5. Reactions were conducted at 37 °C for 30 min before.

### Preparation of duramycin and 4 (non-enzymatic reaction) *in vitro*

Compound **1** was incubated with MBP-DurN (5:1, peptide: enzyme) in 100 mM Tris-HCl pH 7.5, 300 mM KCl, and 10 mM MgCl2 at 37 °C for 12 h. No DTT addition to the peptide indicated full conversion of **1**. The reaction mixture was purified by analytical HPLC (see General Methods) and fractions containing **1** (judged by MS) were lyophilized. The dried peptide was dissolved in 50 mM Tris pH 8.5, then aminopeptidase was added at 0.1 U/μg of peptide, and the reaction mixture was incubated at 37 °C for 12 h to remove the residual leader peptide residues. The reaction mixture was analyzed by analytical HPLC (General Methods), MALDI-TOF-MS, LC-MS, and LC/MSMS indicating production of duramycin. For the preparation of **4**, compound **1** was incubated in 50 mM Tris-HCl pH 10 at r.t. for 12 h and the DTT assay indicated full consumption of **1**. Purification and removal of residual leader peptide residues to yield **4** was carried out as described above. Peptide concentrations were estimated by comparing the peak area at 214 nm in the LC chromatogram to that of authentic duramycin.

### Activity assays for DurN and its mutants with 1 and its variants

All assays were performed with 50 μM modified DurA or its variants with 10 μM or 5 μM MBP-DurN or its mutants in 100 mM Tris-HCl pH 7.5, 300 mM KCl, and 50 mM MgCl2. A positive control with MBPDurN and compound **1** and a negative control with just MBP and compound **1** were included as well as a negative control where MBP-DurN (or its mutants) was omitted; control reactions were conducted in parallel to the reaction of interest under the same conditions. Reactions were incubated at 37 °C for 1 h then treated with DTT using the procedure described above. Samples were analyzed by LC-MS and/or LC-MS/MS and relative activities measured by end-point assays. The area of peaks of products with and without DTT adduct were determined, and reaction yield was calculated as [area of non-adduct]/([area of non-adduct]+[area of adduct]).

### Protein crystallization

Pooled fractions from MBP tag removal and purification of MBP-DurN (see Supplementary Information 1) were concentrated to ≤ 5 mL and injected onto a 120 mL Superdex 200 10/300 GL column (GE Healthcare) to purify by SEC using 300 mM KCl, 20 mM HEPES pH 7.5 as running buffer.

*DurN (apo)*: Following SEC, DurN was concentrated to 7 mg/mL and crystallized in 2 µL hanging drops at 16 °C whereby DurN solution was mixed 1:1 (*v:v*) with a reservoir solution containing 25% polyethylene glycol (PEG) 3350, 0.2 M magnesium chloride, and 0.1 M bis-tris pH 6.5. Apo DurN also produced similar diffraction quality crystals using a different reservoir solution containing 20% PEG 8000, 0.2 M magnesium acetate, and 0.1 M sodium cacodylate pH 6.0. In both cases, rod-shaped crystals grew to a maximum size within 48 h.

*DurN + compound **1**-Dha6Ala*: Following SEC, DurN was concentrated to 9 mg/mL and incubated with 1.8 mM compound **1**-Dha6Ala for 30 min on ice. This solution was mixed 1:1 (*v:v*) with a reservoir solution containing 25% PEG 3350, 0.2 M MgCl2, and 0.1 M bis-tris pH 6.0 in 2 µL hanging drops incubated at 16 °C. Hexagonal shaped crystals grew to maximum size within 48 h. *DurN + duramycin:* Following SEC, DurN was concentrated to 6 mg/mL and incubated with 1.5 mM duramycin for 30 min on ice. Insoluble particulates were pelleted by centrifugation and the supernatant was mixed 1:1 (*v:v*) with a reservoir solution containing 22.5% PEG 3350, and 0.1 M 2-(*N-*morpholino)ethanesulfonic acid (MES) pH 5.6 in 2 µL hanging drops at 16 °C. Thick rod-shaped crystals grew to a maximum size within 72 h.

### Crystallographic data collection, structure solutions, and refinement

Data were collected at the Advanced Photon Source at Argonne National Lab using the Life-Science Collaborative Access Team 21-ID-D, 21-ID-F, and 21-ID-G beamlines. Before collecting data, crystals were cryoprotected by immersing the crystals in similar solutions supplemented with 20% ethylene glycol and then flash-freezing in liquid nitrogen. To obtain phases experimentally, co-crystals of DurN and compound **1**-Dha6Ala were soaked with 0.5 M potassium iodide for 24 h at 16 °C and used to collect diffraction data at 8 keV. Data were initially processed using autoPROC^23^ and further processed in autoSHARP to obtain phase information. The initial model was rebuilt using Buccaneer^24^ and refined using REFMAC5^25^ in the CCP4 Software Suite. Diffraction datasets corresponding to apo DurN crystals and DurN-duramycin co-crystals were phased after similar processing in autoPROC followed by molecular replacement using Phaser MR^26^. Ligand parameters for compound **1**-Dha6Ala and duramycin were determined using Phenix eLBOW^27^ and modeled into their corresponding electron densities, followed by iterative rounds of automated REFMAC5 refinement and manual rebuilding in Coot^28^. Solvent molecules and potassium ions were first incorporated using Phenix Refine^29^ or REFMAC5 then manually curated in Coot.

### Preparation and purification of compound Dur-FL

Duramycin (1 mg) was dissolved in 50 mM HEPES pH 7.5. Carboxyfluorescein succinimidyl ester (2 mg) was dissolved in 10 μL of dimethyl sulfoxide and added (total reaction volume = 1 mL). The reaction was left at r.t. in the dark for 2 h, desalted and analyzed by MS indicating both doubly and singly acylated duramycin. The products were purified by analytical HPLC (see General Methods), lyophilized, and stored at −20 °C. Peptide concentration was estimated by comparing the peak area at 214 nm to that of duramycin and NHS-Fluorescence standard.

### Fluorescence polarization (FP) binding studies

Serial dilutions (1:1) for MBP-DurN or its mutants were prepared with FP buffer (300 mM KCl, 20 mM HEPES, pH 7.5) in a 96-well PCR plate. Each well contained 45 μL solution. The highest MBP-DurN concentration was 15 μM. For each well, 45 µL Dur-FL of a 20 nM stock solution was added, to give a final concentration of 10 nM. Then, 80 μL from each well was transferred to a well in a black 384-well plate (i.e. each well contains 80 μL liquid, with 15 uM˜500 pM MBP-DurN or its mutants, and 10 nM Dur-FL). FP analysis was performed on a BioTek Synergy H4 Hybrid reader. The detection method was set to “fluorescence” and read type was “endpoint”. Measurements were made by excitation at 485 nm (20 nm bandwidth) and emission at 528 nm (20 nm bandwidth). The Corning^TM^ black 384-well plate (Corning^TM^ 3575) was incubated in the plate reader for 5 min at 25 °C, shaken for 1 min, and followed by data collection. Polarization was calculated as Polarization = [I_(parallel)_-I_(perpendicular)_]/[I_(parallel)_-I_(perpendicular)_]. Dissociation constant (K_d_) values were calculated using a dose-response curve (equation below) in Origin Pro 9.1 (OriginLab) with three independent titrations^30^. Background fluorescence from the proteins alone was subtracted from the fluorescence polarization signal obtained with the fluorophore.

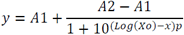

Where *y* is polarization, *A1* is the minimum polarization, *A2* is the maximum polarization, *X0* is the IC50, *p* is the Hill coefficient, and *x* is the concentration of Dur-FL.

### Competition fluorescence polarization (FP) binding studies

The same procedures were applied for all competition FP studies unless specified otherwise. The inhibitor of interest was prepared at 200 μM in water, and serially diluted 1:1 with FP buffer (300 mM KCl, 20 mM HEPES, pH 7.5) to provide 16 wells with concentrations ranging from 100 µM to 0.003 µM in a 96-well PCR plate. A solution of MBP-DurN (2 µM) and Dur-FL (20 nM) was prepared in the same FP buffer and incubated at r.t. for 5 min before transferring 45 µL to each well, then 80 μL of the mixtures in each well were transferred to a black-bottomed 384-well plate (i.e. 80 μL wells containing 1 μM MBPDurN, 10 nM Dur-FL, and inhibitor concentration ranging from 100 μM to 3 nM). IC50 values were calculated from the 50% saturation point using a dose-response curve fit in Origin Pro 9.1 (OriginLab) with three independent titrations^30^. All IC50 values were transferred to Ki values by applying the Munson-Rodbard equation (see equation below)^31^.

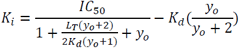

Where *yo* is the initial bound to free ratio for the labeled species before perturbation of equilibrium by the added inhibitor, and *LT* is the total amount of labeled species.

### Derivatization of lysinoalanine (Lal) standard

MeOD-d4 (3 mL) was cooled in a 50 mL pear-shaped glass flask immersed in an ice-water bath. Acetyl chloride (0.9 mL) was added dropwise. This solution was added to 2 mg Lal standard (LL and DL-lysinoalanine mixture, Bachem) and the reaction mixture was heated at 110 °C for 1 h under reflux, then allowed to cool down to r.t. in air. The solvent was removed by a stream of nitrogen gas. Dichloromethane (3 mL) and pentafluoropropionic anhydride (1 mL) were added to the flask cooled in an ice-water bath. The mixture was heated at 110 °C under reflux for 30 min, allowed to cool to r.t., and the solvent was removed under a stream of nitrogen gas for 2 h. For each sample, 800 μL of pyridine was used to dissolve all resulting compounds, and the solutions were directly analyzed by GC-MS.

### Hydrolysis and derivatization of duramycin, compound 4, and deoxyduramycin

The same hydrolysis and derivatization procedures were applied for all Lal-containing samples. To 1 mg of peptide, 3 mL of 6 M deuterium chloride in D2O was added to the sample (˜1 mg) in a glass pressure tube (Ace Glass). The reaction mixture was heated without stirring at 80 °C for 4 h. The solution was transferred to a 50 mL pear-shaped bottle, and the solvent was removed with a rotary evaporator. Then the same derivatizing procedures were used as described for the derivatization of Lal standard.

### GC-MS analysis of lysinoalanine standard, duramycin, compound 4, and deoxyduramycin

The derivatized samples were analyzed by GC-MS using an Agilent 7890 gas chromatograph equipped with a ZB-1MS column (Phenomenex, 30 m × 0.25 mm × 0.25 μm). Samples were dissolved in 300 L of pyridine and a small volume introduced to the instrument via a splitless injection at a flow rate of 1.6 mL/min helium gas. The front inlet was set to 230 °C for initial temperature, the purge flow was 20 mL/min, the purge time was 1 min, and the total flow was 24 mL/min. The flow rate in the column was 1.6 mL/min. The temperature method used for the oven was 100 °C for 2 min, followed by 100 °C to 215 °C at 2.5 °C /min, and 215 °C for 2 min. The mass spectrometer was operated in simultaneous scan/selected-ion monitoring (SIM) mode, monitoring at the characteristic fragment masses of 190 Da, 230 Da, 405 Da, 420 Da, 433 Da, 468 Da, 643 Da, and 705 Da, for Lal residues.

### MD simulations

Unconstrained MD simulations in water were performed using the Amber 16 and AmberTools 17 package^32^. Parameters for dehydro amino acids, (methyl)lanthionine, Hya and Lal were generated using the general AMBER force field (*gaff2*)^32^, with partial charges set to fit the electrostatic potential generated at the HF/6-31G(d) level by the RESP (restrained electrostatic potential) model^33^. Water molecules were treated with the SHAKE algorithm. Long-range electrostatic effects were modelled using the particle mesh Ewald method^34^. An 8 Å cutoff was applied to Lennard–Jones and electrostatic interactions. The Andersen equilibration scheme was used to control and equalize the temperature with a 2 fs time step at a constant volume and temperature of 300 K. Production trajectories were run for 1.0 μs.

## References

1. Kessler H. et al. The structure of the polycyclic nonadecapeptide Ro 09-0198. Helv. Chim. Acta 71, 1924-1929 (1988).

2. Hayashi F. et al. The structure of PA48009: the revised structure of duramycin. J. Antibiot. 43, 1421-1430 (1990).

3. Fredenhagen A. et al. Duramycins B and C, two new lanthionine containing antibiotics as inhibitors of phospholipase A2. Structural revision of duramycin and cinnamycin. J. Antibiot. 43, 1403-1412 (1990).

4. Chen E. et al. Mathermycin, a Lantibiotic from the Marine Actinomycete *Marinactinospora thermotolerans* SCSIO 00652. Appl. Environ. Microbiol. 83 (2017).

5. Kodani S., Komaki H., Ishimura S., Hemmi H. & Ohnishi-Kameyama, M. Isolation and structure determination of a new lantibiotic cinnamycin B from *Actinomadura atramentaria* based on genome mining. J. Ind. Microbiol. Biotechnol. 43, 1159-1165 (2016).

6. Smith T.E. et al. Accessing chemical diversity from the uncultivated symbionts of small marine animals. Nat. Chem. Biol. 14, 179-185 (2018).

7. Pomorski T., Hrafnsdóttir S., Devaux P.F. & van Meer G. Lipid distribution and transport across cellular membranes. Semin. Cell Dev. Biol. 12, 139-148 (2001).

8. Iwamoto K. et al. Curvature-dependent recognition of ethanolamine phospholipids by duramycin and cinnamycin. Biophys. J. 93, 1608-1619 (2007).

9. Hosoda K. et al. Structure determination of an immunopotentiator peptide, cinnamycin, complexed with lysophosphatidylethanolamine by ^1^H-NMR. J. Biochem. 119, 226-230 (1996).

10. Richard A.S. et al. Virion-associated phosphatidylethanolamine promotes TIM1-mediated infection by Ebola, dengue, and West Nile viruses. Proc. Natl. Acad. Sci. U.S.A. 112, 14682-14687 (2015).

11. Oliynyk I., Varelogianni G., Roomans G.M. & Johannesson, M. Effect of duramycin on chloride transport and intracellular calcium concentration in cystic fibrosis and non-cystic fibrosis epithelia. Apmis 118, 982-990 (2010).

12. Zhao M., Li Z. & Bugenhagen, S. 99mTc-labeled duramycin as a novel phosphatidylethanolamine-binding molecular probe. J. Nucl. Med. 49, 1345-1352 (2008).

13. Delvaeye T. et al. Noninvasive whole-body imaging of phosphatidylethanolamine as a cell death marker using (99m)Tc-duramycin during TNF-induced SIRS. J. Nucl. Med. (2018).

14. Huo L., Ökesli A., Zhao M. & van der Donk W.A. Insights into the Biosynthesis of Duramycin. Appl. Environ. Microbiol. 83, e02698-02616 (2017).

15. Ökesli A., Cooper L.E., Fogle E.J. & van der Donk W.A. Nine post-translational modifications during the biosynthesis of cinnamycin. J. Am. Chem. Soc. 133, 13753-13760 (2011).

16. Burley S.K. et al. RCSB Protein Data Bank: Sustaining a living digital data resource that enables breakthroughs in scientific research and biomedical education. Protein Sci. 27, 316-330 (2018).

17. Holm L. & Rosenström, P. Dali server: conservation mapping in 3D. Nucleic Acids Res. 38, W545 (2010).

18. Vagin A.A. et al. REFMAC5 dictionary: organization of prior chemical knowledge and guidelines for its use. Acta Crystallogr. D Biol. Crystallogr. 60, 2184-2195 (2004).

19. Tang W., Jiménez-Osés G., Houk K.N. & van der Donk W.A. Substrate control in stereoselective lanthionine biosynthesis. Nat. Chem. 7, 57-64 (2015).

20. Lopatniuk M., Myronovskyi M. & Luzhetskyy, A. *Streptomyces albus*: A new cell factory for non-canonical amino acids incorporation into ribosomally synthesized natural products. ACS Chem. Biol. 12, 2362-2370 (2017).

21. O’Rourke S., Widdick D. & Bibb, M. A novel mechanism of immunity controls the onset of cinnamycin biosynthesis in *Streptomyces cinnamoneus* DSM 40646. J. Ind. Microbiol. Biotechnol. 44, 563-572 (2017).

22. Gibson D.G. et al. Enzymatic assembly of DNA molecules up to several hundred kilobases. Nat. Methods 6, 343-345 (2009).

23. Vonrhein C. et al. Data processing and analysis with the autoPROC toolbox. Acta Crystallogr. D Biol. Crystallogr. 67, 293-302 (2011).

24. Cowtan K. The Buccaneer software for automated model building. 1. Tracing protein chains. Acta Crystallogr. D Biol. Crystallogr. 62, 1002-1011 (2006).

25. Murshudov G. et al. REFMAC5 for the refinement of macromolecular crystal structures. Acta Crystallogr. D Biol. Crystallogr. 67, 355-367 (2011).

26. McCoy A.J. et al. Phaser crystallographic software. J. Appl. Crystallogr. 40, 658-674 (2007).

27. Moriarty, N.W., Grosse-Kunstleve, R.W. & Adams, P.D. electronic Ligand Builder and Optimization Workbench (eLBOW): a tool for ligand coordinate and restraint generation. Acta Crystallogr. D Biol. Crystallogr. 65, 1074-1080 (2009).

28. Emsley P., Lohkamp B., Scott W.G. & Cowtan, K. Features and development of Coot. Acta Crystallogr. D Biol. Crystallogr. 66, 486-501 (2010).

29. Afonine P.V. et al. Towards automated crystallographic structure refinement with phenix.refine. Acta Crystallogr. D Biol. Crystallogr. 68, 352-367 (2012).

30. Zhang Z. et al. Biosynthetic Timing and Substrate Specificity for the Thiopeptide Thiomuracin. J. Am. Chem. Soc. 138, 15511-15514 (2016).

31. Munson, P.J. & Rodbard, D. An Exact Correction to the “Cheng-Prusoff” Correction. J. Recept. Res. 8, 533-546 (1988).

32. Wang J., Wolf R.M., Caldwell J.W., Kollman P.A. & Case, D.A. Development and testing of a general amber force field. J. Comput. Chem. 25, 1157-1174 (2004).

33. Bayly C.I., Cieplak P., Cornell W.D. & Kollman, P.A. A well-behaved electrostatic potential based method using charge restraints for deriving atomic charges: the RESP model. J. Phys. Chem. 97, 10269-10280 (1993).

34. Darden T., York D. & Pedersen, L. Particle mesh Ewald: An *N*-log(N) method for Ewald sums in large systems. J. Chem. Phys. 98, 10089-10092 (1993).

